# A switch–rheostat circuit governs quorum sensing homeostasis in a phytopathogen

**DOI:** 10.64898/2026.07.23.740290

**Authors:** Nemanja Ristović, Iris Bertani, Gianluca Triolo, Michael P. Myers, Cristina Bez, Vittorio Venturi

## Abstract

*Pseudomonas fuscovaginae*, a wide-host-range plant pathogen of several cereal and grass species, possesses two canonical *N*-acyl homoserine (AHL)-based quorum sensing (QS) systems called PfsI/R and PfvI/R, both of which are inactive under laboratory conditions but active in planta. The *pfsI*-*pfsR* intergenic region encodes for RsaM, a putative protein that has since been hypothesized to act as a repressor switch of this QS circuit. In the present study, we demonstrate that the stringent repression exerted on the PfsI/R system depends entirely on the divergent promoter/intergenic *pfsR-rsaM* region rather than RsaM itself. Remarkably, this regulatory switch element stringently represses the expression of both the *pfsR* and *rsaM* genes. We further show for the first time that RsaM is endogenously expressed and functions as a negative regulator modulating the PfsI/R circuit instead of preventing its activation. Taken together, our results evidence a unique two-tiered/hierarchical repression of a QS system, provided by a master repressor switch and a repressor modulator.

**Importance:** *Pseudomonas fuscovaginae* is a globally occurring plant pathogen that employs AHL QS to regulate virulence. In this bacterium, QS signaling circuits display a rather unusual feature: the lack of activation at high cell densities under standard laboratory conditions. A hypothetical regulator named RsaM was previously linked to this phenomenon as a repressor switch acting on the PfsI/R AHL QS system in the absence of an unknown signal or stimulus. In this study, we show that a regulatory element within the *pfsR-rsaM* intergenic region acts as the primary switch of the PfsI/R QS system, independently of RsaM. Conversely, RsaM functions as a post-activation modulator that fine-tunes the QS response. This study advances our understanding of the regulatory configurations of unconventional QS systems by revealing a two-tier control mechanism in which a master regulatory switch governs circuit activation, while the previously uncharacterized protein RsaM controls signaling output once the system is engaged.

## Introduction

*Pseudomonas fuscovaginae* is a broad-host-range phytopathogen associated with brown sheath rot of rice and other cereals. It is capable of colonizing various plant niches, both as an endophyte and an epiphyte, and displays considerable plasticity in its life cycle (Rott, 1989; Bigirimana et al., 2015). Several virulence factors and virulence-associated processes, including phytotoxins, secretion systems, and quorum sensing (QS), contribute to its pathogenicity (Batoko et al., 1997; Mattiuzzo et al., 2011; Patel et al., 2014).

QS is a bacterial cell-cell communication process mediated by diffusible signaling molecules that links cell density and environmental stimuli to transcriptional regulation of target gene expression, enabling coordination of various social behaviors. Among Pseudomonadota (formerly Proteobacteria), the most common QS systems consist of signaling and sensing modules, comprised of a LuxI family synthase, the producer of signaling molecules *N*-acyl homoserine lactones (AHLs), and a LuxR family transcriptional regulator, the receptor of cognate AHLs. QS typically relies on low basal level of constitutive production of AHLs, which accumulate as cell density increases. Once critical AHL concentrations are reached, these signals are detected by the LuxRs, which then dimerize and bind the *lux* boxes, the *cis*-regulatory elements located within the promoter regions of LuxR target genes. One of the key regulatory targets of the LuxRs-AHL is the cognate *luxI* gene, setting off an autoinducing loop. This results in signal amplification, gene expression synchronization across the bacterial population, and a shift from individual to group behaviors. LuxR regulons are diverse, encompassing various genes involved in the production of common goods, and thus enabling coordination of distinct collective endeavors (Miller & Bassler, 2001; Whitehead et al., 2001; Waters & Bassler, 2005). AHL-based QS systems in bacterial phytopathogens often regulate phenotypes involved in pathogenicity, such as motility, virulence factor production, biofilm formation, and oxidative stress tolerance (Quiñones et al., 2005; Hussain et al., 2008; Põllumaa et al., 2012).

The timing and magnitude of QS activation usually do not align only with cell density but are also affected by extracellular and intracellular cues (Kindler et al., 2019; Striednig & Hilbi, 2022; Schuster et al., 2023). These signals are conveyed via protein and RNA regulators that act most commonly at the transcriptional, or, more rarely, at the post-transcriptional or post-translational level, and affect the amount or activity of the LuxR and LuxI proteins. While some regulators of the LuxI/R systems function globally and link QS to other regulatory cascades, others feature as intrinsic/built-in elements, typically counteracting the LuxR-driven positive feedback loop. QS is thus today regarded as a regulatory network that integrates a diverse array of cues beyond cell density (Venturi et al., 2011; Frederix & Downie, 2011; Spacapan et al., 2023).

*P. fuscovaginae* possesses two AHL-based QS modules, named PfsI/R (producing and responding to several types of unsubstituted AHLs) and PfvI/R (producing and responding to several types of 3-oxo-substituted AHLs). Strikingly, neither system exhibits conventional activation dynamics under standard laboratory conditions, remaining shut down at high cell densities. Importantly, the knockout mutants of the synthase or regulator genes display impaired virulence in rice (*Oryza sativa*) and quinoa (*Chenopodium quinoa*), providing evidence that AHL QS in *P. fuscovaginae* is activated during plant infection (Mattiuzzo et al., 2011). These findings indicated involvement of additional cues and regulatory pathways which, combined with AHL concentration, unlock one or both QS systems. In a previous genetic screen of a Tn*5* genomic mutant library of *P. fuscovaginae* UPB0736, a mutant was identified having a dramatic increase in *pfsI* promoter activity and overproducing PfsI-synthesized AHLs. The Tn*5* mutant was hypothesized to be inserted into a novel gene designated as *rsaM* which is intergenically located between *pfsI* and *pfsR*. Two possible open reading frames (ORFs) encoding RsaM, which are in frame with one another, are possible in this intergenic region (Fig. 1A) (Mattiuzzo et al., 2011). It was therefore hypothesized that RsaM could be a stringent repressor of the PfsI/R system that is likely to act as a switch in response to a yet unknown signal/s. The sequence of RsaM is unique, with no significant homology to any characterized proteins. Similar ORFs have subsequently been identified in a number of species; interestingly, the putative *rsaM* gene was always located adjacent to AHL QS loci (Venturi et al., 2023). RsaM homologue from *Burkholderia cenocepacia*, BcRsaM, was structurally characterized, evidencing that it possesses a distinct domain architecture. Several sequence- and domain-organization features appear to be maintained across the so-far predicted RsaMs, with the hallmark being the presence of a preserved tryptophan quartet (Michalska et al., 2014; Venturi et al., 2023). While all these findings suggested that *rsaM* ORFs are endogenously expressed, no functional studies have been performed to date. The BcRsaM lacks known DNA- or RNA-binding motifs, enzyme active site signatures, and is unable to bind AHLs (Michalska et al., 2014). AHL production in *B. thailandensis*, *B. glumae*, and *Acinetobacter baumannii* is overactivated upon disruption of *rsaM* homologs, suggesting that putative RsaMs invariably feature as negative regulators of QS circuits (Le Guillouzer et al., 2018; López-Martín et al., 2021; Goo & Hwang, 2023). As *rsaM* mutants in several bacterial species exhibit decreased virulence, this repression has also been linked to maintenance of QS homeostasis (Mattiuzzo et al., 2011; Chen et al., 2012; López-Martín et al., 2021).

**Figure 1.**
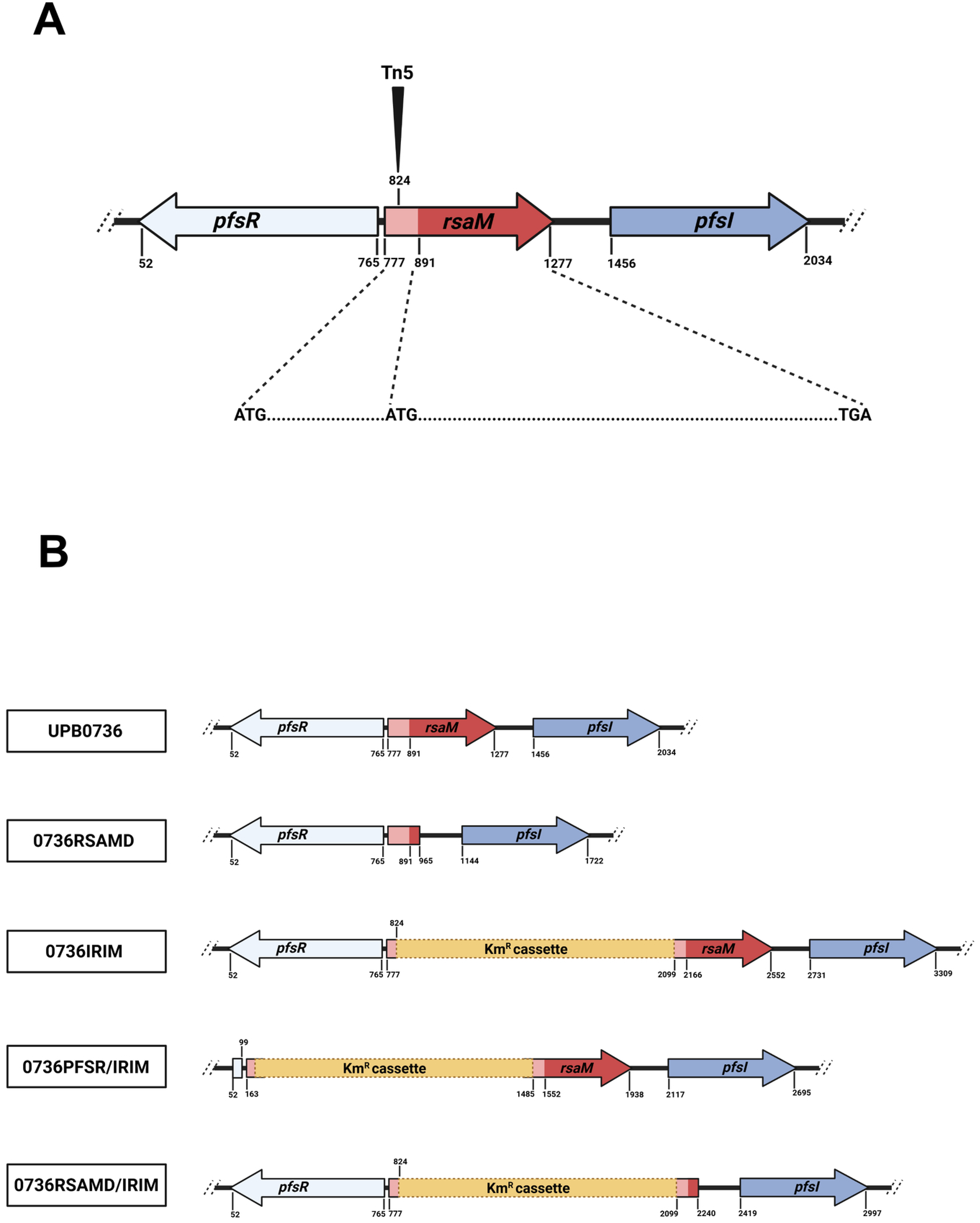
Genomic architecture of the *pfsR/rsaM/pfsI* locus. A) The 2150 bp region in *P. fuscovaginae* UPB0736 (wild-type strain) containing *pfsR*, *pfsI* and *rsaM* genes. The shorter ORF of the *rsaM* gene is shown in dark red. Long *rsaM* ORF encompasses the same region as well as an upstream sequence (light red), thus encoding a 38 aa longer protein. The position of the Tn*5* insertion leading to hyperactivation of the PfsI/R system is also indicated. B) Genomic organization of the *pfsR*/*rsaM*/*pfsI* locus in *P. fuscovaginae* UPB0736 (wild-type strain) and mutant derivatives generated in this study: *P. fuscovaginae* 0736RSAMD (in-frame deletion of the *rsaM* gene in the wild-type background), *P. fuscovaginae* 0736IRIM (kanamycin resistance cassette inserted at the position of the previously described Tn*5* insertion), *P. fuscovaginae* 0736PFSR/IRIM (in-frame deletion of the *pfsR* gene in the mutant harboring the kanamycin resistance cassette), and *P. fuscovaginae* 0736RSAMD/IRIM (in-frame deletion of the *rsaM* gene in the mutant harboring the kanamycin resistance cassette).

The transposon insertion within the *pfsR-pfsI* intergenic region in *P. fuscovaginae* UPB0736 resulting in PfsI-AHL hyperproduction was mapped to the possible *N*-terminus-coding region of the longer *rsaM* ORF (Fig. 1A), suggesting, as mentioned above, that this phenotype could be due to the interruption of the coding sequence of the *rsaM* gene (Mattiuzzo et al., 2011). In this study, we now evidence that the shorter ORF of the *rsaM* gene is the functional ORF, and demonstrate that the *pfsR-rsaM* intergenic region contains a pivotal genetic switch element of the PfsI/R system that stringently controls both the *pfsR* and *rsaM* gene expression. We then evidence that RsaM is not a master repressor of the PfsI/R system as first postulated, but functions as a modulator that attenuates this regulatory circuit once the genetic switch is in an ON state. The findings presented here support a model in which the PfsI/R QS system of *P. fuscovaginae* is controlled by two hierarchical layers of repression. The first consists of a regulatory switch located within the *pfsR–rsaM* intergenic region that maintains the circuit in an OFF state. The second involves RsaM, which does not prevent activation of the system but instead acts after the circuit has switched ON to negatively modulate QS output, thereby maintaining signaling homeostasis. This two-layered regulatory architecture suggests that *P. fuscovaginae* maintains its QS system in a repressed, ecologically silent state until specific environmental or host-associated cues permit activation, allowing virulence-associated signaling to emerge only under appropriate conditions.

## Materials and methods

### Bacterial strains and growth conditions

*E. coli* strains were grown in Luria-Bertani (LB) medium (tryptone 10 g/l, yeast extract 5 g/l, NaCl 10 g/l) supplemented with antibiotics when necessary (ampicillin 100 mg/l, kanamycin 100 mg/l, gentamicin 10 mg/l, tetracycline 10 mg/l) at 37 °C with shaking. For overexpression of the recombinant RsaM, *E. coli* M15 carrying the plasmids pQE31RsaM and pREP-4 was grown in Terrific broth (TB) medium (tryptone 12 g/l, yeast extract 24 g/l, glycerol 4 g/l, KH_2_PO_4_ 2.3 g/l, K_2_HPO_4_ 12.5 g/l) with added ampicillin 100 mg/l and kanamycin 25 mg/l. *Pseudomonas fuscovaginae* UPB0736 and mutant derivatives were grown in King’s B (KB) medium (proteose peptone no.3 20 g/l, MgSO_4_ x 7H_2_O 1.5 g/l, KH_2_PO_4_ 1.2 g/l, glycerol 10 g/l) at 30° C with shaking. When required, the medium was supplemented with antibiotics (kanamycin 100 mg/l, gentamicin 40 mg/l, tetracycline 40 mg/l, nitrofurantoin 100 mg/l). To make solid media, 13 g/l agar was added.

### Plasmid construction/recombinant DNA techniques

The plasmids and primers used in this study are listed in Table 1 and Table 2, respectively. Routine DNA manipulation techniques, including restriction enzyme digestion, agarose gel electrophoresis, DNA fragment purification, ligation with T4 ligase, PCR, and transformation into *E.coli*, were performed as described previously (Sambrook, 1989).

**Table 1.** Plasmids used in this study.

| Plasmid | Characteristics | Reference or source |
| --- | --- | --- |
| pGEM-T | Cloning vector; Amp<sup>r</sup> | Promega |
| pQE31 | Expression vector; ColE1; Amp<sup>r</sup> | Qiagen GmbH, Hilden, Germany |
| pLAH30 | Expression vector; IncP; Tc<sup>r</sup> | (Bertani et al., 1999) |
| pMP220 | Promoter probe vector; IncP; Tc<sup>r</sup> | (Spaink et al., 1987) |
| **pEX19Gm** | Suicide vector for targeted mutagenesis, Gm<sup>r</sup> | (Hmelo et al., 2015) |
| **pBBR1MCS-5** | Broad-host-range vector, Gm<sup>r</sup> | (Kovach et al., 1995) |
| **pRK2013** | Km<sup>r</sup> Tra<sup>+</sup> Mob<sup>+</sup> ColEI | (Figurski & Helinski, 1979) |
| **pUC4K** | pUC7 derivative, Amp<sup>r</sup> and Km<sup>r</sup> | Addgene, Watertown, MA |
| **pREP-4** | Contains *lacI*, Km<sup>r</sup>, p15A replicon | Qiagen GmbH, Hilden, Germany |
| **pJRLACI** | pJRD215 containing *lacI*, Km<sup>r</sup> | (Bertani et al., 1999) |
| **pEX19Gm_A** | pEX19Gm containing construct for the generation of *P. fuscovaginae* 0736RSAMD | This study |
| **pEX19Gm_B** | pEX19Gm containing construct for the generation of *P. fuscovaginae* 0736IRIM | This study |
| **pEX19Gm_E** | pEX19Gm containing construct for the generation of *P. fuscovaginae* 0736RSAMD/IRIM | This study |
| **pEX19Gm_J** | pEX19Gm containing construct for the generation of *P. fuscovaginae* 0736PFSR/IRIM | This study |
| **pUC57_B** | pUC57 containing construct for generating pEX19Gm_B | This study |
| **pPFSI220** | pMP220 with the *pfsI* promoter cloned upstream of the *lacZ* gene | This study |
| **pRSAM220** | pMP220 with the putative short *rsaM* ORF promoter region cloned upstream of the *lacZ* gene | This study |
| **pRSAML220** | pMP220 with the putative long *rsaM* ORF promoter region cloned upstream of the *lacZ* gene | This study |
| **pPFSR220** | pMP220 with the *pfsR* promoter cloned upstream of the *lacZ* gene | This study |
| **pBBRSAM** | pBBR1MCS-5 with the short ORF of the *rsaM* gene under control of the *lac* promoter | This study |
| **pBBRSAML** | pBBR1MCS-5 with the long ORF of the *rsaM* gene under control of the *lac* promoter | This study |
| **pBBRSAMN** | pBBR1MCS-5 with the short *rsaM* ORF with nonsense mutation of start codon under control of the *lac* promoter | This study |
| **pBBRSAMnat** | pBBR1MCS-5 with the *rsaM* gene under control of its native promoter | This study |
| **pBBPFSR** | pBBR1MCS-5 with the *pfsR* gene under control of the *lac* promoter | This study |
| **pQE31RsaM** | pQE31 with the *rsaM* gene cloned in frame with the His-tag-coding region of the vector | This study |
| **pLAH30RsaM** | pLAH30 with the *rsaM* gene cloned in frame with the His-tag-coding region of the vector | This study |
| **pLAH30PfsR** | pLAH30 with the *pfsR* gene clone in frame with the His-tag-coding region of the vector | This study |

**Table 2.**
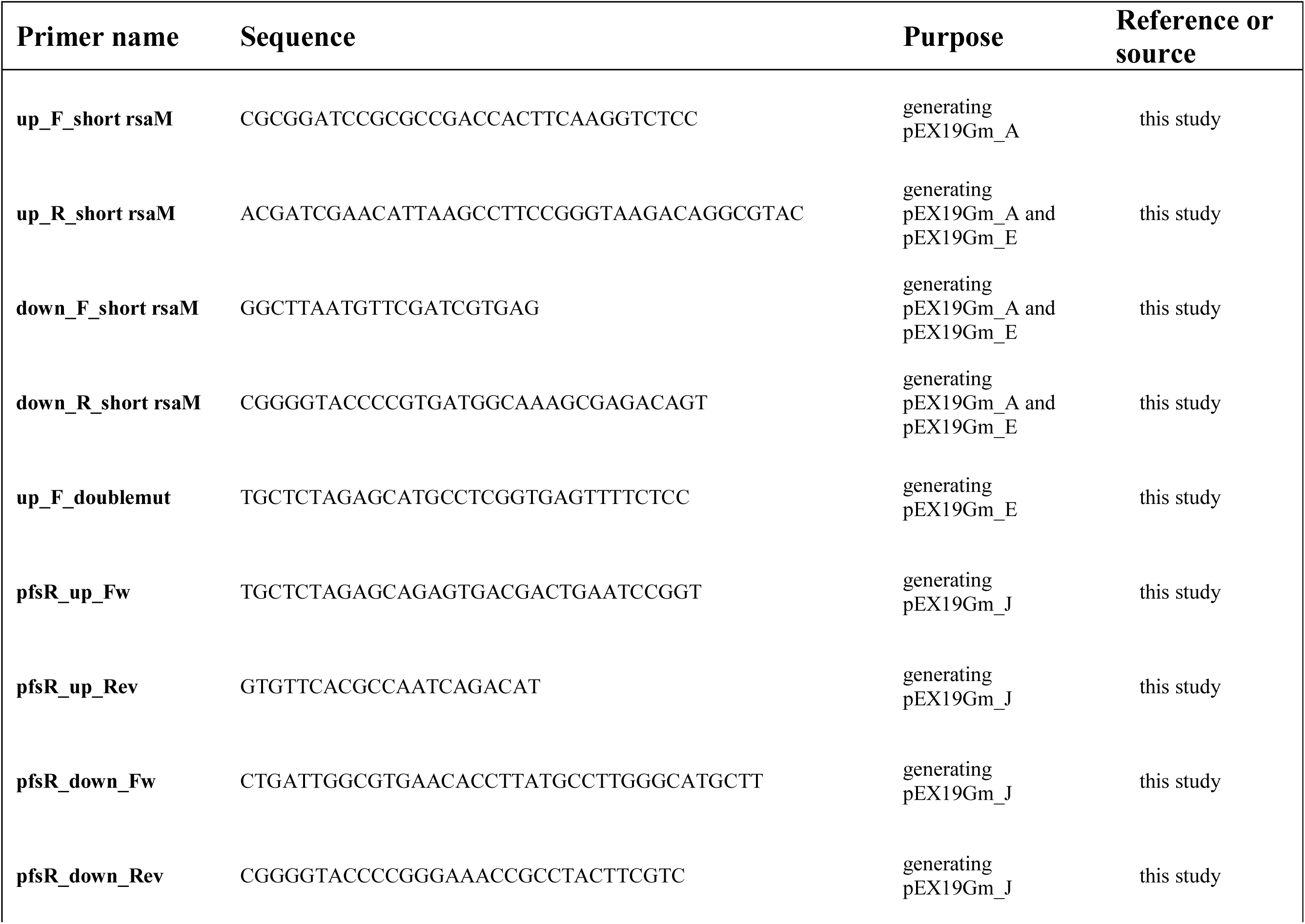
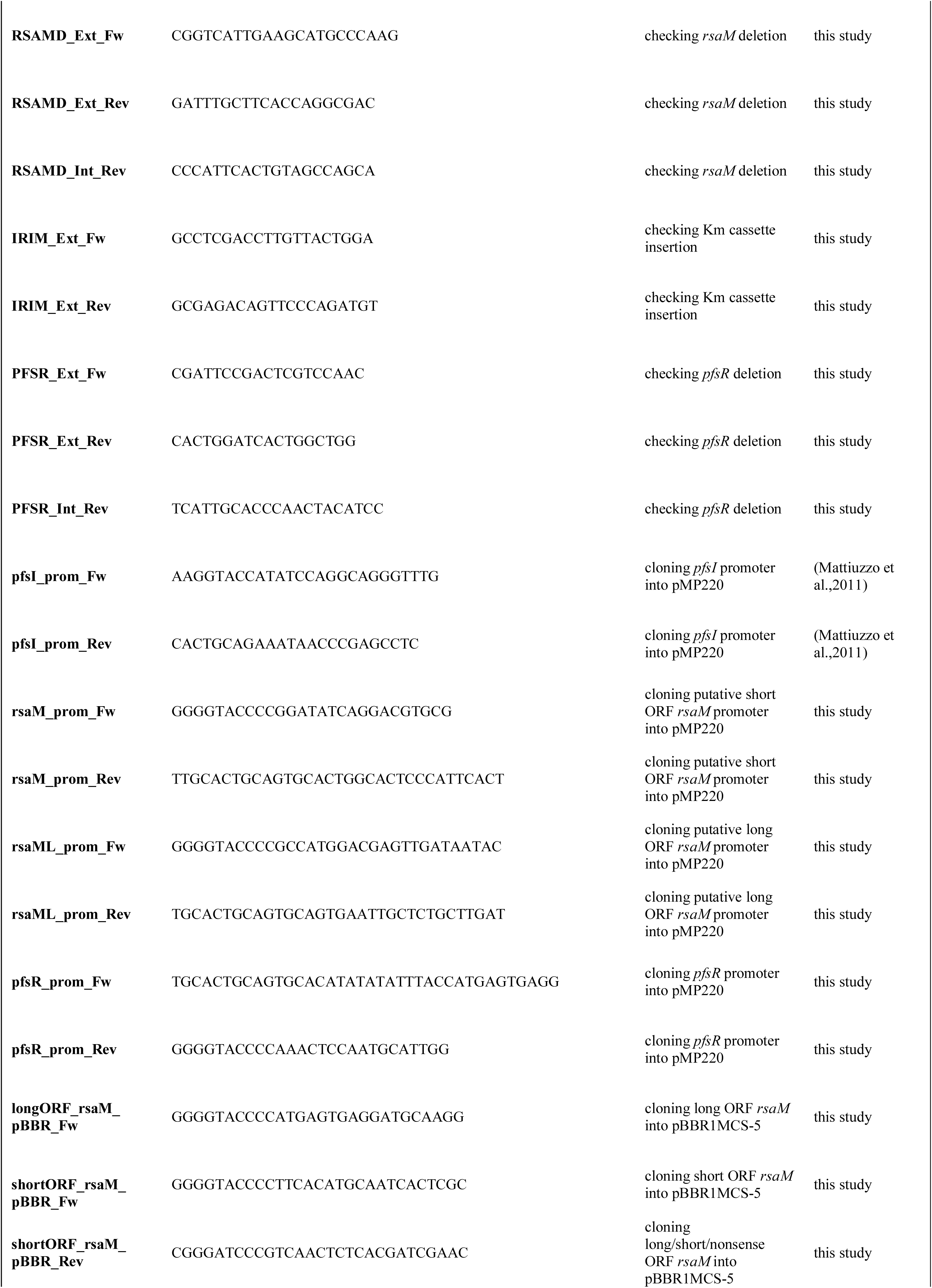
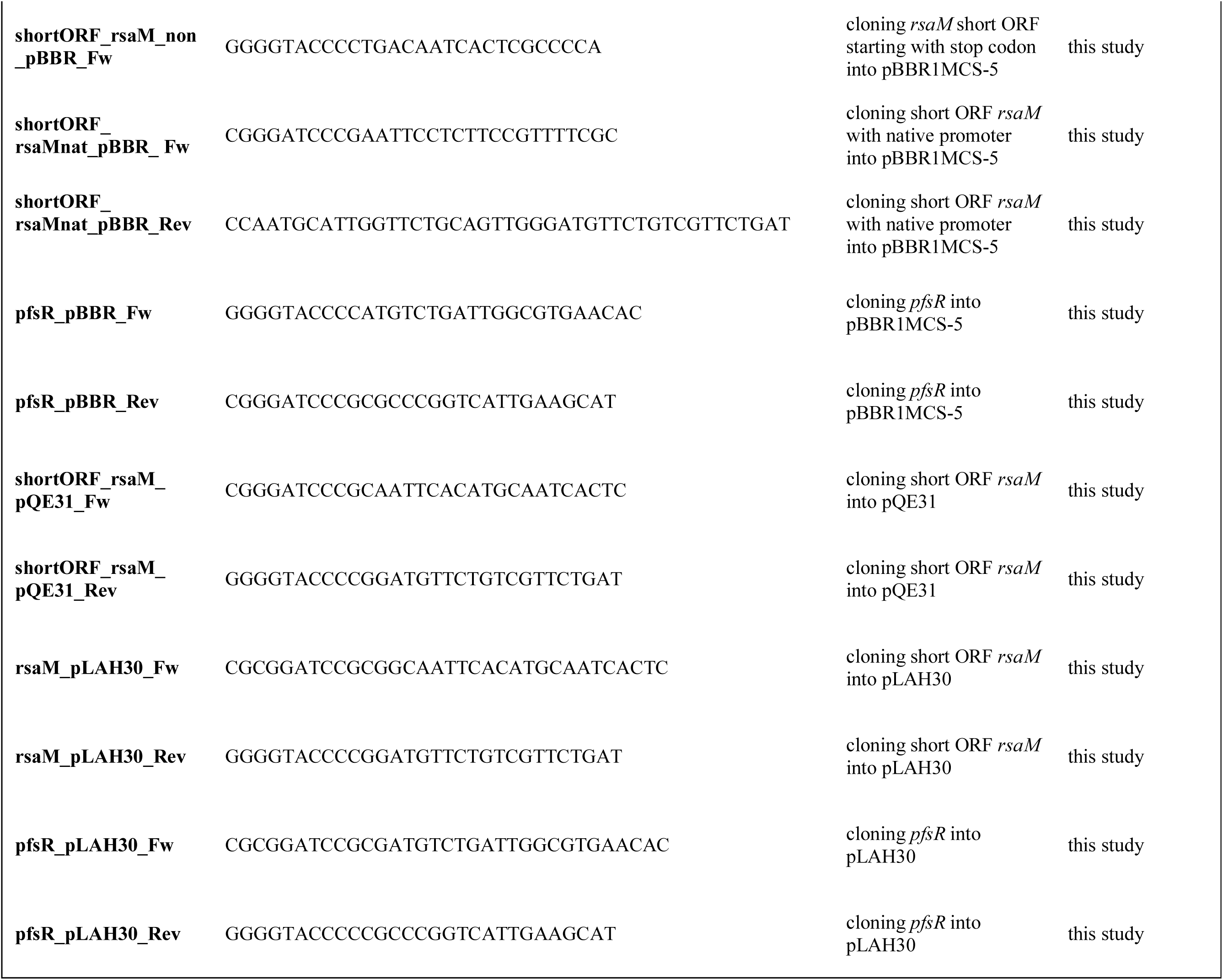
Primers used in this study.

Generation of *P. fuscovaginae* UPB0736 mutants

Targeted mutagenesis was performed using the suicide vector pEX19Gm, via two-step allelic exchange, as described previously (Hmelo et al., 2015). To generate the *rsaM* deletion mutant in the wild-type background (*P. fuscovaginae* 0736RSAMD), 700 bp upstream and 500 bp downstream of the sequence targeted for removal were first PCR amplified and fused via overlap extension PCR. The fused product was then cloned into pEX19Gm, yielding pEX19Gm_A. To obtain the mutant with the kanamycin cassette inserted in the position of the previously described Tn*5* insertion (*P. fuscovaginae* 0736IRIM), the pUC57 plasmid containing 500 bp upstream and downstream of the target position separated by the BamHI restriction site was synthesized by Genescript. This 1000 bp fragment was cut from the vector and cloned into pEX19Gm. The obtained plasmid was then linearized with BamHI and ligated to the Km cassette excised from the pUC4K plasmid with the same enzyme, thus generating the construct pEX19Gm_B. To obtain the double mutant having both the kanamycin resistance cassette insertion and *rsaM* deletion (*P. fuscovaginae* 0736RSAMD/IRIM), 500 bp upstream and downstream of the target sequence were amplified via PCR using genomic DNA of 0736IRIM as a template. Overlap extension PCR was used to generate the fusion of the two amplicons, which was then cloned into pEX19Gm to yield pEX19Gm_E. Similar strategy was employed to obtain the construct pEX19Gm_J, which was subsequently used to generate the *pfsR* deletion in 0736IRIM (mutant 0736PFSR/IRIM). All constructs were conjugated from *E.coli* DH5α into *P. fuscovaginae* UPB0736 (for mutants 0736RSAMD and 0736IRIM) or *P. fuscovaginae* 0736IRIM (for the mutants 0736RSAMD/IRIM and 0736PFSR/IRIM) by triparental mating on solid KB with *E. coli* HB101 harboring the plasmid pRK2013. Conjugation mixtures were incubated overnight at 30°C before transfer onto KB plates containing 50 mg/l nitrofurantoin and 30 mg/l gentamicin to select for transconjugants. The second recombination event was induced by plating putative transconjugants onto NSLB (LB medium without added NaCl) supplemented with 100 mg/l nitrofurantoin and 15% (m/V) sucrose. Colonies were screened for the presence of deletions or insertions by PCR, using primers with binding sites outside of the region cloned into pEX19Gm. The mutants were confirmed by PCR and sequencing. The genomic maps of the *pfsI/R* locus in *P. fuscovaginae* UPB0736 and mutant strains 0736RSAMD, 0736IRIM, 0736PFSR/IRIM, and 0736RSAMD/IRIM are shown in Figure 1B.

### β-galactosidase promoter activity assays

To assess the activities of the *pfsI*, *pfsR*, and *rsaM* promoters, the corresponding regions were amplified using primer pairs given in Table 2 and genomic DNA of *P. fuscovaginae* UPB0736 as template, followed by cloning into the transcriptional reporter vector pMP220. The constructs were checked for correct sequence and introduced into the wild-type and mutant strains via triparental mating. For the β-galactosidase assay, test strains were grown for 24 h in 5 ml of KB medium in flasks. The exception was the test in which the mutant 0736RSAMD harbored constitutively expressed PfsR, which was performed after 16 h. The tests were conducted essentially as described previously (Miller, 1972), with modifications (Stachel et al., 1985). Negative controls, with test strains carrying empty pMP220, were included in each test. All promoter activity values (expressed in Miller units) shown in the graphs are the averages of three biological replicates, with error bars indicating standard deviation.

### AHLs detection and extraction followed by HPLC/MS-MS

As the PfsI synthase produces exclusively unsubstituted AHLs (Mattiuzzo et al., 2011), the extent of activation of the PfsI/R circuit in various genetic backgrounds was qualitatively assessed via T-streaks with the AHL biosensor *Chromobacterium violaceum* 026 (McClean et al., 1997), which does not respond to the major AHLs produced by the PfvI synthase (3-oxo-C10 and 3-oxo-C12) (Morohoshi et al., 2008). A single colony of each test strain was streaked in line perpendicular to the biosensor on KB plates. Plates were incubated for 72 h at 30 °C.

In order to validate that the *C.violaceum* 026 is a reliable biosensor for PfsI-AHLs, extraction and relative quantification of AHLs from spent supernatants of liquid cultures of the wild-type *P. fuscovaginae* UPB0736 and the major mutant derivatives within the *pfsI-rsaM-pfsR* region was performed. To match the conditions on the biosensor plates, the liquid cultures were grown until late stationary phase (24 h) in KB medium (5 ml, in 50 ml Falcon tubes). The spent supernatants were obtained by centrifuging 1 ml of each culture at 8000 rcf for 5 min, followed by filtration using 0.22 µm filters. Caffeine (the internal standard) was added to the supernatants to a final concentration of 25 nM, and formic acid to a final concentration of 21 mM. Water-saturated ethyl acetate was added to the acidified supernatants in a 1:1 ratio, followed by mixing in an eppendorf mixer for 5 min. The mixture was then centrifuged (max speed, 5 min), and the organic phase was removed and left to evaporate at room temperature for 3 h. The samples were stored at -20 °C until analysis. Extracts were reconstituted into 100 µL of 0.1% formic acid in water, and 10 µL of the reconstituted extract was injected onto a 100 µm × 20 cm column packed with Intersustain AQ 3 µm beads (GL Sciences) and developed with a 5%–80% gradient of Acetonitrile in 0.1% formic acid in 20 min. The effluent of the column was directed into the orifice of a 6550 QTOF (Agilent). The QTOF was run in targeted ion mode with quantitation based on the conversion of the precursor mass to the 102.055 *m*/*z* product ion. The elution times of the AHLs were scouted using purified standards, and caffeine was added to each sample and detected by the transition from 194.99 *m/z* to 138.066 *m/z*. The peaks were extracted, and the peak areas were determined using the MSHunter software package (Agilent). AHLs were quantified according to their peak area, relative to the peak area of added caffeine. This ratio was subsequently normalized to OD_600_ for each AHL. Mean summed peak area ratio/OD_600_ for unsubstituted and 3-oxo-substituted AHLs is depicted in the graphs. The extraction and analysis were performed for four biological replicates.

### RNA isolation and sequencing

Cultures of each strain (five biological replicates) were grown in 5 ml of KB medium in flasks until the onset of the stationary phase. 1000 µl of RNAprotect Bacteria Reagent (Qiagen) was added to 500 µl of culture (containing approximately 10^9^ CFU), vortexed for 5 s, and centrifuged at 2500 rcf at 4 °C for 10 min. The supernatant was thoroughly removed, and cell pellets were immediately transferred to dry ice and stored at -80 °C until use. Total RNA was isolated using the RNeasy Mini Kit (Qiagen) following the manufacturer’s instructions with a few modifications. Namely, cells were lysed in TE buffer (Tris 10 mM, EDTA 1 mM, pH 8) with the addition of 5 mg/ml of lysozyme, followed by 20 s of vortexing and 10 min incubation at room temperature. RNA concentration and purity were assessed with Nanodrop, while RNA integrity was verified on a 1.5 % (m/V) agarose gel. RNA sequencing using the Illumina NovaSeq X Plus Sequencing System, as well as the subsequent bioinformatic and statistical analysis, was conducted by Novogene. The sequencing data were deposited in NCBI Bioproject under the accession number PRJNA1370452.

### Recombinant protein expression, purification and anti-RsaM antibodies generation

An overnight culture of *E. coli* M15 harboring the plasmids pREP-4 and pQE31RsaM was diluted in 250 ml fresh TB medium (starting OD_600_ 0.05). Upon reaching the mid-exponential phase (OD_600_ 0.6), IPTG was added to the culture (final concentration 1 mM) which was then incubated for 5 h (37 °C, with shaking). The culture was centrifuged at 2500 rcf at 4 °C for 25 min, and the pellets were stored at 4 °C until purification. Since the recombinant RsaM localized entirely in the insoluble fraction of the lysate, a protocol for the purification of inclusion bodies was used prior to affinity purification. Briefly, the cells were resuspended in 1 mM dithiothreitol (DTT) and disrupted using a French press (2.5 kbar, 4 °C), followed by centrifugation of the lysate at 9300 rcf at 4 °C for 30 min. The pellet was washed two times in a buffer (50 mM Tris-HCl buffer, 5 mM DTT, 5 mM EDTA, 0.75 % (m/V) sodium deoxycholate, pH 8) and once in water, each time followed by centrifugation at 9300 rcf at 4 °C for 30 min. The purified inclusion body was solubilized in Tris-urea buffer (10 mM Tris, 8 M urea, 100 mM NaH_2_PO_4,_ pH 8), and the recombinant RsaM was purified under denaturing conditions using the QIAexpressionist kit (Qiagen), according to the manufacturer’s instructions. Bradford reagent (Bio-Rad) was used to determine protein concentration. The expression of the recombinant protein and its presence throughout purification were confirmed via standard SDS-PAGE, as described previously (Laemmli, 1970). Anti-RsaM antibodies were raised in rabbits and affinity-purified at ThermoFisher Scientific.

### Western blot and IP-MS

To obtain samples for Western blots, 1 ml of liquid cultures (grown 4 h, 16 h or 24 h) was centrifuged at 3300 rcf for 5 min, followed by resuspension of the cell pellets in 10 cell volumes of sample buffer (4 % (m/V) SDS, 20 mM DTT, 0.01% (m/V) bromophenol blue, 60 mM Tris, pH 8), boiling (95 °C, 5 min), sonication (30 sec at full force), and centrifugation at 9300 rcf for 10 min. Equal volumes of cleared lysate (7.5 µl) were loaded onto a discontinuous SDS-PAGE gel (5% stacking, 12% resolving). Separated proteins were transferred onto a PVDF membrane activated with PBST buffer with added methanol (1xPBS, 0.1% (V/V) Tween, 10% (V/V) methanol) via wet transfer. The membrane was then incubated in a PBST buffer containing 5% (m/V) skimmed milk (2 h at room temperature), followed by incubation with anti-RsaM antibodies (1:500 dilution in PBST) overnight at 4 °C. The membrane was then washed three times with PBST buffer (7 min each wash) and incubated with a 1:2500 dilution of the anti-IgG-rabbit HRP-conjugated secondary antibody in PBST for 1 h at room temperature. After three washes with PBST, the membrane was incubated with ECL (Bio-Rad), as described by the manufacturer. Pictures of the membrane were taken on ChemiDoc (Bio-Rad). For immunoprecipitation, 1 ml of the overnight liquid cultures (24 h) was centrifuged (2500 rcf, 4 °C, 5 min). The pellets were washed once in PBS, resuspended in 1 ml of the RIPA buffer (150 mM NaCl, 1% (V/V) NP-40, 0.5% (m/V) sodium deoxycholate, 0.1% (m/V) SDS, 50 mM Tris, pH 8), followed by sonication (on ice, 20 s at maximum intensity, four times with 1 min pauses). The lysate was cleared by centrifugation at 10000 rcf at 4 °C for 10 min, passed through a 0.22 µm filter, and incubated with the affinity-purified anti-RsaM antibodies (final concentration 500 µg/ml) for 1 h at 4 °C. The EZview^TM^ Protein G Affinity Gel (Sigma) was then added to the lysate, with the rest of the immunoprecipitation protocol performed according to the manufacturer’s instructions. For MS, samples were digested directly on the beads by the addition of 100 ng of trypsin in 20 μl of 20 mM triethylammonium bicarbonate (pH 8.5) for 16 h at room temperature. Following digestion, the supernatant was desalted using STAGE tips fabricated from Empore Octadecyl C18 disks (Sigma-Aldrich). The samples were aspirated from the beads and passed two times over a STAGE tip using air pressure provided by a 10 ml syringe (Eppendorf) and immediately eluted with 15 μl of 65% acetonitrile in 0.1% formic acid. The samples were dried and resuspended in 10 μl of 0.1% formic acid and injected onto a custom-fabricated 250 mm × 0.075 mm fused silica column. The column was custom-packed using 2.8 μm Ascentis Express RPA resin (Sigma-Aldrich). The column was developed over 60 min using a gradient of 0.1% formic acid to 80% acetonitrile using a 1290 Infinity 2 UHPLC (Agilent). The gradient ran from 3% ACN to 27% ACN in the first 50 minutes, then to 80% ACN in the next 5 minutes and followed by a further 5 minutes at 80% ACN. The effluent of the column was sprayed directly into a 6550 QTOF (Agilent) using a custom-built ESI nanospray source. Analysis was performed with the precursor mass range set to 325 to 1700 m/z and each precursor scan was followed by 15 fragmentation scans using a dynamic exclusion window of 6 seconds, an isolation width of 1.5 Da and a fragmentation scan mass range of 50 to 1700 m/z. The resulting spectra were converted into peak lists using Mass Analyst version 10.1 (Agilent), and spectral matching was performed using the X!Tandem search engine (Fury version). The databases were automatically concatenated by X!Tandem along with a list of common contaminants. Spectral matching was performed using a precursor mass error of +/- 25 ppm and a fragmentation error of +/- 25 ppm.

### Statistical analysis

Welch’s T-tests were used to compare the means obtained in promoter assays. The mean relative levels of AHLs produced by different strains were analyzed using Welch’s one-way ANOVA, followed by the post hoc Dunnett T3 multiple comparisons test. The analysis was performed in Prism 11.0.2 (GraphPad Software).

## Results

### RsaM does not maintain the OFF state of the PfsI/R quorum sensing circuit

We hypothesized that activation of the PfsI/R system in the previously reported *rsaM*::Tn*5* mutant (Mattiuzzo et al., 2011) was either due to the interruption of the *rsaM* coding sequence or the *rsaM* promoter. The *rsaM* gene has two possible ORFs in the same frame, one having a 38 amino acid extension at the *N*-terminus (Fig. 1A). It was therefore initially of interest to confirm the presence of RsaM in the wild-type *P. fuscovaginae* UPB0736 and elucidate the correct ORF. To do this, we cloned the shorter ORF of *rsaM* into an expression vector and successfully expressed the His-tagged protein in *E. coli*. The recombinant protein was purified and used to raise polyclonal antibodies as described in the Materials and Methods section. The antibodies were then used in Western blot analysis and immunoprecipitation coupled with mass spectrometry (IP-MS). Surprisingly, RsaM was not detected in the wild-type strain with either technique (data not shown). In line with this, putative promoters of the *rsaM* gene showed very low activity (putative promoter for the short *rsaM* ORF) or no activity (putative promoter for the long *rsaM* ORF) (Fig. S1) while the transcriptomic analysis peformed here evidenced an extremely low level of transcription from the *rsaM* locus (median DESeq2-normalized expression count less than 10) (Fig. 2A). It was thus concluded that *rsaM* is transcribed at very low levels in the wild-type strain, while its putative protein product was undetectable by the techniques used in this study.

**Figure 2.**
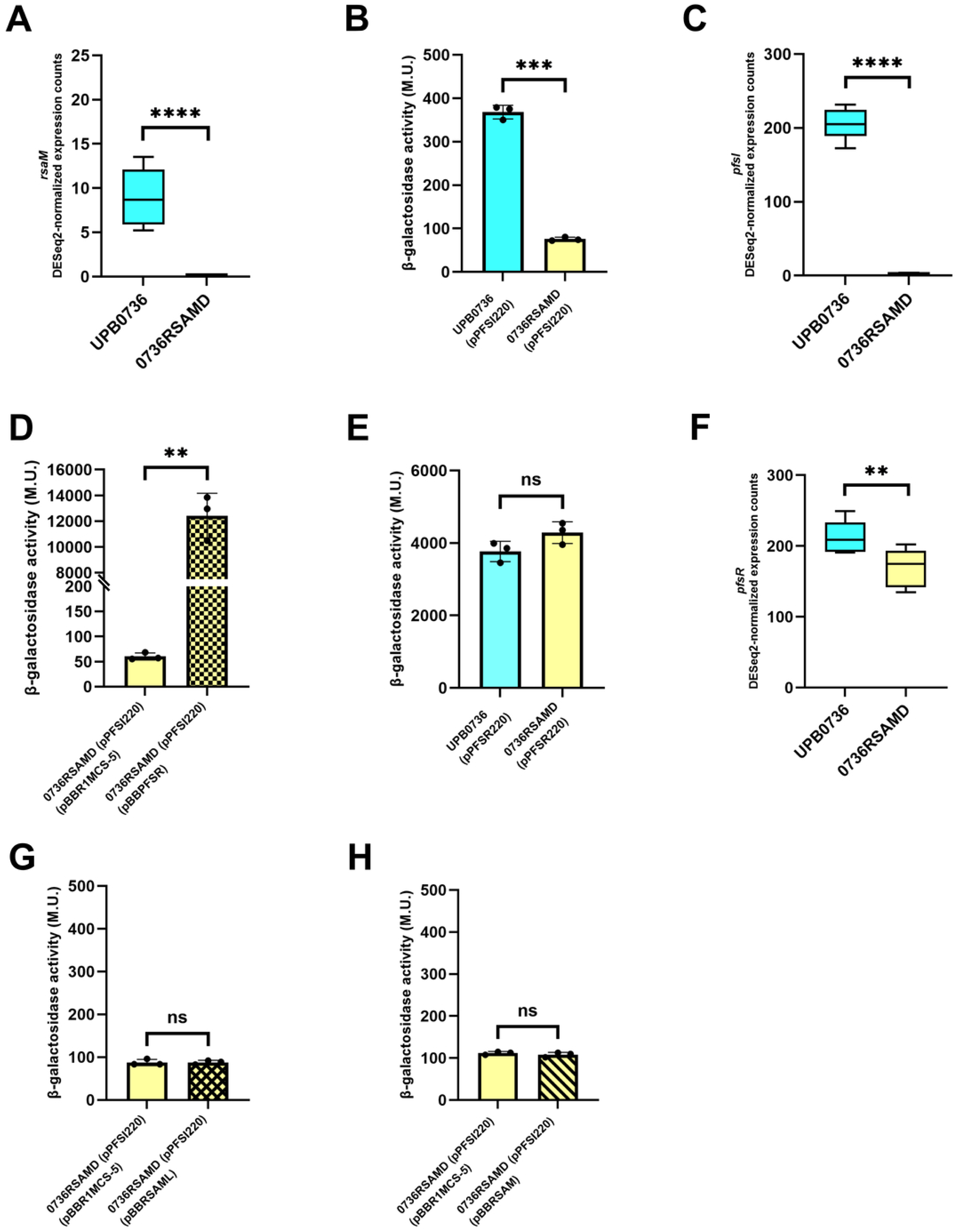
The *rsaM* gene is not required for the maintenance of repression keeping the PfsI/R system silenced. A) DESeq2-normalized counts for *rsaM* transcripts in the wild-type *P. fuscovaginae* UPB0736, obtained through the analysis with the mutant lacking the *rsaM* gene (*P. fuscovaginae* 0736RSAMD). B) *pfsI* promoter activity in *P. fuscovaginae* UPB0736 and 0736RSAMD. C) DESeq2-normalized counts for *pfsI* transcripts in *P. fuscovaginae* UPB0736 and 0736RSAMD. D) *pfsI* promoter activity in *P. fuscovaginae* 0736RSAMD harboring pBBR1MCS-5 or pBBPFSR (*pfsR* under control of the *lac* promoter). E) *pfsR* promoter activity in *P. fuscovaginae* UPB0736 and 0736RSAMD. F) DESeq2-normalized counts for *pfsR* transcripts in *P. fuscovaginae* UPB0736 and 0736RSAMD. G) *pfsI* promoter activity in *P. fuscovaginae* 0736RSAMD harboring pBBR1MCS-5 or pBBRSAML (*rsaM* long ORF under control of the *lac* promoter). H) *pfsI* promoter activity in *P. fuscovaginae* 0736RSAMD harboring pBBR1MCS-5 or pBBRSAM (*rsaM* shorter ORF under control of the *lac* promoter). The promoter activity data are presented as means of three biological replicates ± standard deviation. Asterisks indicate significant differences between the means (*p<0.05; **p<0.01; ***p<0.001; ****p<0.0001). All promoter assays were performed with negative controls (test strains carrying empty pMP220); the corresponding β-galactosidase activities are listed in Table S1. The DESeq2-normalized read counts are presented with box plots that depict the medians (central lines), the interquartile ranges (boxes), and minimum and maximum data values (whiskers) for each set of samples (five biological replicates for each strain). Asterisks indicate significant differences between two groups of samples (*p<0.05; **p<0.01; ***p<0.001; ****p<0.0001).

To determine whether loss of function of the *rsaM* gene causes PfsI/R overactivation, targeted mutagenesis was performed to generate an in-frame deletion mutant where most of the shorter *rsaM* ORF (and thus also the long ORF) was deleted (Fig. 1B). Surprisingly, the resulting mutant, *P. fuscovaginae* 0736RSAMD, did not produce AHLs, displayed no *pfsI* promoter activity, and lacked *pfsI* transcripts. This was in contrast with the wild type where very low levels of PfsI-AHLs and *pfsI* transcription were present (Fig. S2 and S3; Fig. 2B and C). To confirm that the deletion of the *rsaM* gene caused no *cis*-acting changes to *pfsI* expression, we constitutively expressed PfsR *in trans* in this mutant. This resulted in the activation of the *pfsI* promoter, evidencing that *pfsI* transcription remained functional (Fig. 2D). The *pfsR* transcript level decreased only slightly in the *rsaM* mutant compared to the wild type; therefore, we concluded that the cessation of *pfsI* transcription was not due to changes in PfsR levels (Fig. 2E and F). We were not able to complement the phenotype of this mutant upon introduction of the long or short *rsaM* ORF under control of a constitutively active *lac* promoter *in trans*; the reason for this is currently unknown (Fig. 2G and H). Taken together, these results indicated that the *rsaM* gene is not transcribed in the wild-type *P. fuscovaginae* UPB0736 under the laboratory conditions and that this locus does not exert a top-level repressive effect on the PfsI/R QS system.

### A master regulatory switch controls PfsI/R quorum sensing

As we established that repression of the PfsI/R circuit is not the direct effect of RsaM, we hypothesized that the *rsaM-pfsR* noncoding intergenic region plays an important role in activating the system. To validate this phenotype in the *rsaM*::Tn*5* mutant (Fig. 1A), we regenerated this mutant by introducing a kanamycin resistance cassette in the exact genomic position of the Tn*5* integration (Fig. 1B). The resulting mutant, *P. fuscovaginae* 0736IRIM, displayed a similar phenotype, having considerably elevated production of PfsI-AHLs (approximately a 250-fold increase) (Fig. S2A and S4) as well as significantly elevated *pfsI* and *pfsR* promoter activities (approximately 16.7x and 1.7x, respectively) and levels of corresponding RNA transcripts as determined from the RNAseq data (19.4x and 3.2x, respectively) compared to the wild-type *P. fuscovaginae* UPB0736 (Fig. 3A, B, C and D). Remarkably, the transcription of the *rsaM* gene in this mutant was also significantly increased compared to the wild type, as evidenced by promoter studies (a 6.3-fold increase in *rsaM* promoter activity) and transcriptomic analysis (a 115-fold increase in *rsaM* gene expression) (Fig. 3E and F). Consistent with this, performing both Western blot analysis and IP-MS showed that the mutant expressed the RsaM protein corresponding to the predicted shorter ORF-encoded form (Fig. 3G and H, Fig. S5). To our knowledge, this is the first evidence of a protein belonging to the RsaM family being natively expressed.

**Figure 3.**
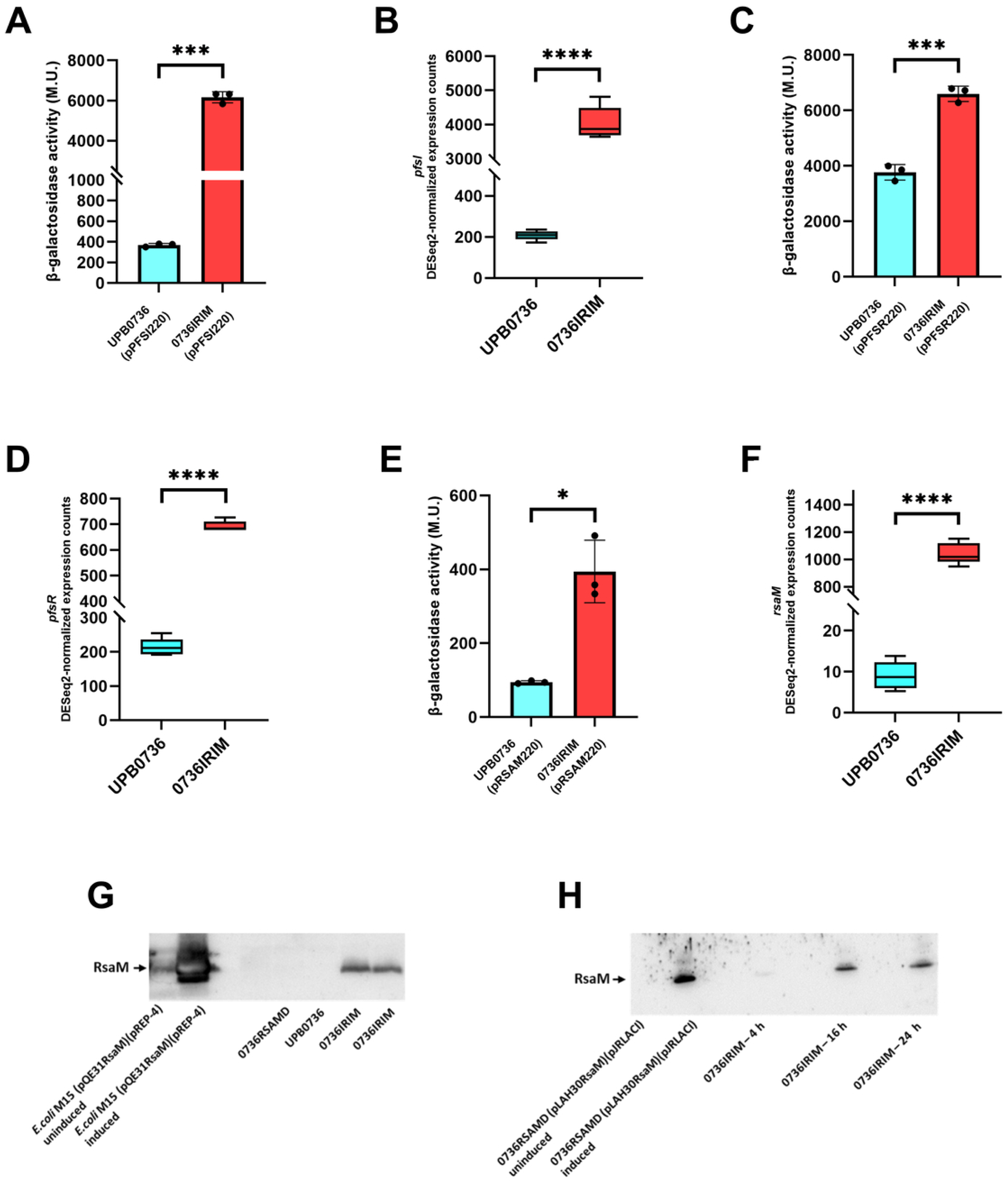
The intergenic *pfsR*-*rsaM* region is crucial in repressing the *pfsR*, *rsaM*, and *pfsI* genes. A) *pfsI* promoter activity in *P. fuscovaginae* UPB0736 and 0736IRIM. B) DESeq2-normalized counts for *pfsI* transcripts in *P. fuscovaginae* UPB0736 and 0736IRIM. C) *pfsR* promoter activity in *P. fuscovaginae* UPB0736 and 0736IRIM. D) DESeq2-normalized counts for *pfsR* transcripts in *P. fuscovaginae* UPB0736 and 0736IRIM. E) *rsaM* promoter activity (short *rsaM* ORF promoter) in *P. fuscovaginae* UPB0736 and 0736IRIM. F) DESeq2-normalized counts for *rsaM* transcripts in *P. fuscovaginae* UPB0736 and 0736IRIM. G) Western blots performed with anti-RsaM antibodies on the total lysates of *E. coli* M15 (pREP-4) harboring pQE31RsaM without (lane 1) or with induction with IPTG (lane 2), *P. fuscovaginae* 0736RSAMD (lane 5), wild-type *P. fuscovaginae* UPB0736 (lane 6), and *P. fuscovaginae* 0736IRIM (lanes 7 and 8). H) Western blots performed with anti-RsaM antibodies on the total lysates of *P. fuscovaginae* 0736RSAMD (pJRLACI) harboring pLAH30RsaM without (lane 1) or with added IPTG (lane 2), *P. fuscovaginae* 0736IRIM in late exponential phase (lane 5), mid-stationary phase (lane 8), and late stationary phase (lane 11). Lysates of the wild-type strain and *P. fuscovaginae* 0736RSAMD obtained at the same time points are in lanes 3, 6, 9, and 4, 7, 10, respectively. Full pictures of Western blots are shown in the Supplementary (Fig. S14). The promoter activity data are presented as means of three biological replicates ± standard deviation. Asterisks indicate significant differences between the means (*p<0.05; **p<0.01; ***p<0.001; ****p<0.0001). All promoter assays were performed with negative controls (test strains carrying empty pMP220); the corresponding β-galactosidase activities are listed in Table S1. The DESeq2-normalized read counts are presented with box plots that depict the medians (central lines), the interquartile ranges (boxes), and minimum and maximum data values (whiskers) for each set of samples (five biological replicates for each strain). Asterisks indicate significant differences between two groups of samples (*p<0.05; **p<0.01; ***p<0.001; ****p<0.0001).

These findings suggested that the region disrupted by the kanamycin cassette contains, or is the target of, a regulatory switch element that represses expression of both *pfsR* and *rsaM*. To affirm that the activation of the circuit was mediated through PfsR, we constructed an in-frame *pfsR* deletion mutant in the 0736IRIM background (designated 0736PFSR/IRIM) (Fig. 1B). Deletion of *pfsR* completely abolished production of PfsI-derived AHLs (Fig. S6), demonstrating that activation of this QS circuit following disruption of the regulatory switch is strictly dependent on PfsR. These results establish PfsR as the essential downstream effector through which the switch controls QS activation.

Together, our results demonstrated that the QS-activating insertion disrupts a regulatory switch within the divergent *pfsR-rsaM* intergenic region rather than the coding sequence of *rsaM*. This switch is responsible for maintaining the PfsI/R QS system in a strongly repressed OFF state and therefore represents the key determinant of circuit activation. Importantly, *rsaM* is also under the control of this switch. Thus, the same regulatory element simultaneously governs expression of both the PfsI/R module and its downstream modulator, RsaM (see below).

The disruption of the PfsI/R master switch also resulted in the activation of the PfvI/R system, as evidenced by a significant increase in the *pfvR* and *pfvI* transcription (Fig. S7) as well as the production of the PfvI-derived 3-oxo-substituted AHLs in 0736IRIM (Fig. S2B). These results suggest that the switch provides overarching repression on both QS systems in *P. fuscovaginae* UBP0736; whether this is a direct effect or is mediated through activation of the PfsI/R system remains to be investigated and is not the focus of this study.

### RsaM functions as a homeostatic rheostat of the PfsI/R quorum sensing circuit

To establish whether RsaM has any role in regulating the PfsI/R system, we first introduced *rsaM in trans* cloned into a plasmid into the insertional mutant 0736IRIM, either under the control of the native *rsaM* promoter or the *lac* promoter present on the backbone of the vector pBBR1MCS-5. Introduction of the constitutively expressed *rsaM* gene resulted in approximately a 2-fold decrease in *pfsI* promoter activity (Fig. 4A) and a significant decrease in PfsI-AHL production (Fig. S8). Although not statistically significant, we also observed a decrease in *pfsI* promoter activity when the *rsaM* gene was expressed under its own promoter (Fig. 4B). These results suggested RsaM modulates activation of the PfsI/R system when triggered by the disruption of the *pfsR*-*rsaM* intergenic region (see above). To further examine this, we carried out targeted mutagenesis by deleting the *rsaM* gene in the 0736IRIM mutant, thereby attaining activation of the PfsI/R system in the absence of RsaM (Fig. 1B). The obtained double mutant (*P. fuscovaginae* 0736RSAMD/IRIM) produced significantly higher levels of PfsI-derived AHLs compared to the mutant 0736IRIM (approximately a 7.7-fold increase) (Fig. S2A and S9). This effect could be complemented since the production of PfsI-AHLs decreased dramatically upon introduction of the *rsaM* gene under control of the *lac* promoter *in trans* (Fig. S10), as well as *pfsI* promoter activity (Fig. 4C). To verify that the complementation observed here was caused by the RsaM protein, we also generated a construct with *lac* promoter-controlled *rsaM* having a nonsense mutation of the start codon and introduced it into this mutant. The production of PfsI-AHLs was not affected in the presence of this construct (Fig. S11), indicating that trans effects of the *rsaM* under the control of the same promoter are RsaM protein-mediated. Overall, these results evidenced that RsaM functions as an intrinsic negative modulator that limits the activation of the PfsI/R circuit once the system has been triggered. Importantly, while causing a dramatic increase in the level of PfsI-AHLs, the deletion of *rsaM* in 0736IRIM did not result in changes in the production of 3-oxo-substituted PfvI-AHLs, suggesting RsaM modulates exclusively the PfsI/R circuit (Fig. S2B). Interestingly, despite the large difference in the production of PfsI-synthesized AHLs, there was no significant change in *pfsI* promoter activity between the double mutant and its parent mutant strain (Fig. 4D), indicating that RsaM most likely functions post-transcriptionally.

**Figure 4.**
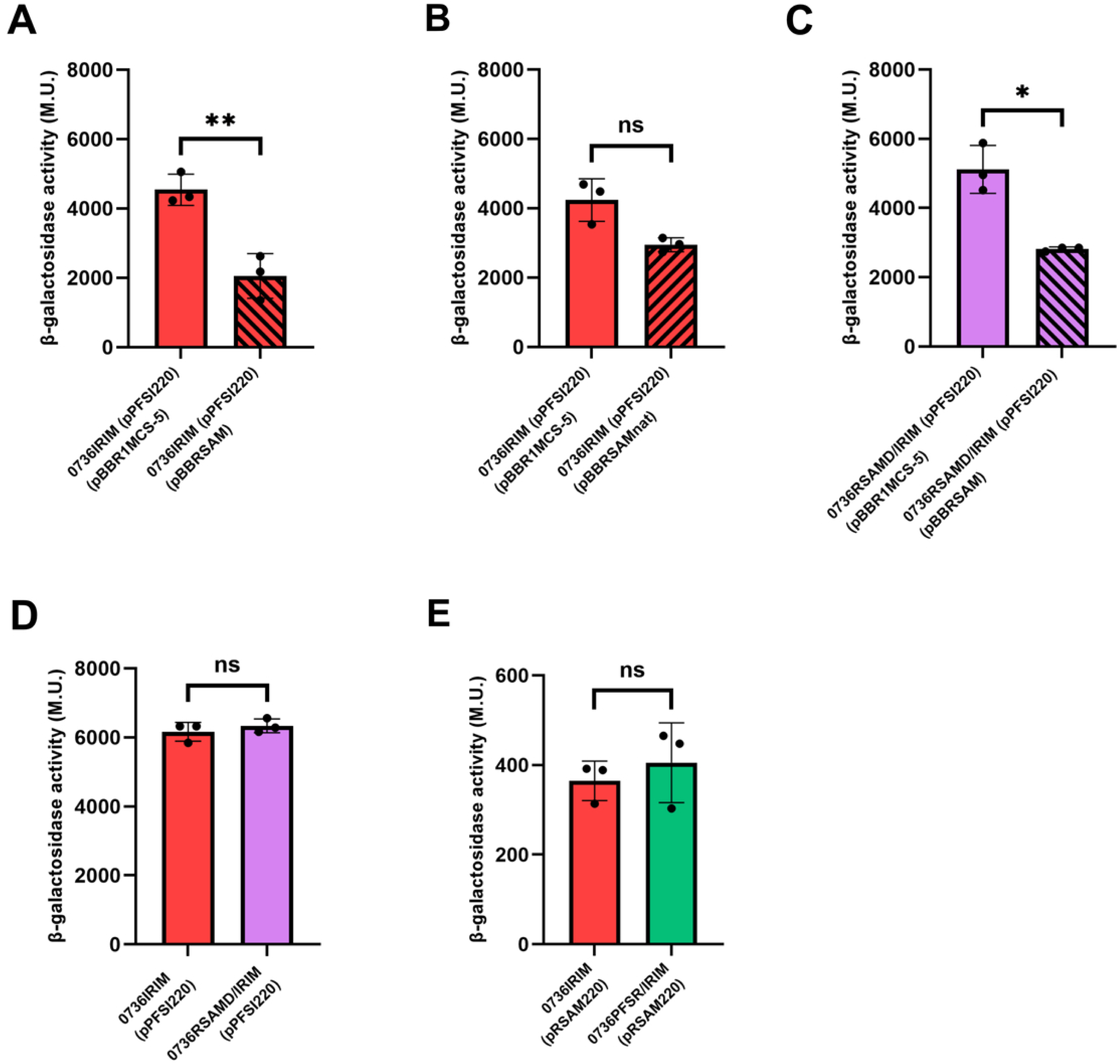
RsaM is a negative modulator of the activated PfsI/R system. A) *pfsI* promoter activity in *P. fuscovaginae* 0736IRIM harbouring pBBR1MCS-5 (control) or pBBRSAM (short ORF *rsaM* under control of the *lac* promoter). B) *pfsI* promoter activity in *P. fuscovaginae* 0736IRIM with pBBR1MCS-5 (control) or pBBRSAMnat (short ORF *rsaM* under control of the native promoter). C) *pfsI* promoter activity in *P. fuscovaginae* 0736RSAMD/IRIM with pBBR1MCS-5 (control) or pBBRSAM (short ORF *rsaM* under control of the *lac* promoter). D) *pfsI* promoter activity in *P. fuscovaginae* 0736IRIM and 0736RSAMD/IRIM. E) *rsaM* promoter activity in *P. fuscovaginae* 0736IRIM and 0736PFSR/IRIM. The promoter activity data are presented as means of three biological replicates ± standard deviation. Asterisks indicate significant differences between the means (*p<0.05; **p<0.01; ***p<0.001; ****p<0.0001). All promoter assays were performed with negative controls (test strains carrying empty pMP220); the corresponding β-galactosidase activities are listed in Table S1.

### The master switch independently controls *pfsR* and *rsaM* expression

To further define the regulatory links within the PfsI/R circuit, we investigated whether *rsaM* is expressed and modulates QS activity in the wild-type strain when the system is forcibly activated by the constitutive expression of *pfsR in trans*. Surprisingly, RsaM could not be detected by Western blot analysis under these conditions, indicating that elevated PfsR levels alone are insufficient to induce *rsaM* expression (data not shown). This observation was unexpected because the wild-type strain constitutively expressing PfsR produced lower amounts of PfsI-AHLs than the *rsaM* deletion mutant (0736RSAMD) carrying the same construct (Fig. S12), suggesting that RsaM-dependent modulation was occurring.

Since endogenous RsaM expression was observed only when the switch in the *pfsR-rsaM* intergenic region was disrupted, we hypothesized that the same regulatory element repressing *pfsR* also controls *rsaM* expression. To examine this, we tested *rsaM* promoter activity in the mutant having both the QS-activating kanamycin cassette insertion in the master switch and the *pfsR* deletion. We found that the activity of the *rsaM* promoter was not affected (Fig. 4E), indicating that the activation of *rsaM* expression does not depend on PfsR and is independently regulated by the master switch.

As the expression of RsaM is blocked by the switch in the wild-type *P. fuscovaginae* UPB0736, it was of interest to affirm the regulatory role of the protein once this top-tier repression is bypassed. To do so, we constitutively expressed *rsaM* from the *lac* promoter in the wild-type background simultaneously expressing *pfsR in trans*, thereby mimicking the regulatory configuration of the 0736IRIM mutant. Under these conditions, RsaM again reduced PfsI/R-dependent AHL production, confirming its function as a negative modulator of the ON-state of this QS circuit (Fig. S13).

Together, these results indicated that the activation of the PfsI/R system and the expression of *rsaM* are coordinated via bidirectional regulatory inputs acting within the *pfsR*-*rsaM* intergenic region, with both depending on deactivation of the same top-tier repressor element.

## Discussion

Phytopathogenic *P. fuscovaginae* possesses two canonical AHL-based quorum sensing (QS) systems that remain silent under standard in vitro growth conditions but become activated in planta and contribute to virulence. A previous Tn*5* insertion in the *pfsI*-*pfsR* intergenic region was shown to strongly activate the PfsI/R system, leading to the hypothesis that the uncharacterized protein RsaM functions as a key repressor of this QS circuit (Mattiuzzo et al., 2011). Our results have demonstrated that rather than disrupting RsaM-mediated repression, the insertion interrupts a previously unrecognized *cis*-regulatory element that acts as a master switch controlling expression of both *pfsR* and *rsaM*. This regulatory element is responsible for maintaining the PfsI/R system in an OFF state under laboratory conditions. In contrast, RsaM does not function as the primary silencer of QS. Instead, once this switch is disrupted and the circuit becomes activated, RsaM acts as a negative feedback regulator that dampens PfsI/R activity and maintains signaling homeostasis. Together, these findings reveal a two-layered regulatory control in which a master switch controls QS activation, while RsaM functions as a homeostatic brake that prevents runaway signaling once the system is engaged.

RsaM was not detected in the wild-type *P. fuscovaginae* under culture conditions used in this study, either via Western blot analysis or IP-MS, and the transcription level of the *rsaM* gene was extremely low. As many bacterial regulators are known to function in a low number of copies per cell, it could not be excluded that this is the case with RsaM as well, with the protein amount being below the detection limit of the techniques used in this study (McAdams & Arkin, 1999; Pulkkinen & Metzler, 2015). However, an in-frame deletion of *rsaM* in the wild-type background did not result in an AHL-overproducing phenotype, suggesting this gene is not pivotal in silencing the PfsI/R circuit. Due to the loss of *pfsI* expression, the mutant did not produce any detectable AHLs. This phenotype and the lack of complementation upon introduction of the intact *rsaM* gene *in trans* raised the possibility that the deletion interrupted a *cis*-regulatory element involved in controlling the transcription of *pfsI*. Interestingly, an in-frame deletion of the *tofM* gene, the *rsaM* homologue in *Burkholderia glumae* 336gr-1, resulted in a small increase in AHL production that could not be complemented by expressing TofM *in trans* (Chen et al., 2012).

Our results evidenced that the *pfsR*-*rsaM* intergenic region has a critical role in repressing the PfsI/R system in the wild-type *P. fuscovaginae*. Activation of the system in the insertional mutant in the intergenic region is triggered by PfsR, which accumulates as a result of increased transcription of the *pfsR* gene. Since the disruption of this region activates the transcription of the *pfsR* gene (thus turning the circuit ON), we have termed the putative regulatory element/regulator acting at this level, as ʺthe switchʺ of the PfsI/R system, the function previously hypothesized to be mediated by RsaM. The same regulatory element is responsible for the stringent repression of the *rsaM* gene in the wild-type strain. Although the exact nature of this regulatory switch element remains to be resolved, it is evident that it acts bidirectionally, repressing the transcription of both the *pfsR* and *rsaM* genes. The large insertion introduced in this study could have impaired the function of the switch by distorting a potential binding site or creating a spacer that impacted the position of the regulator relative to the target promoters (Huo et al., 2009; Levo et al., 2015; Sangeeta et al., 2024). Studies on *rsaM* homologues in *A. baumannii* AB5705 and *B. glumae* BGR1 reported that transposon insertions within the *abaM* and *tofM* genes result in AHL hyperproduction (López-Martín et al., 2021; Goo & Hwang, 2023). Our findings raise the possibility that the hyper-QS phenotypes reported in *A. baumannii* and *B. glumae* may not result from loss of RsaM function, but could also reflect disruption of an upstream regulatory switch. Revisiting these mutants through precise genetic mapping and functional analyses will be important to determine whether this regulatory mechanism is conserved across different bacterial species.

Significantly, deletion of *rsaM* had very different consequences depending on the regulatory state of the QS circuit. The *rsaM* deletion in the wild-type background had little effect, however its deletion in the switch-interrupted strain, in which the PfsI/R system is activated, resulted in strong PfsI-AHL overproduction. Restoration of RsaM expression *in trans* suppressed this phenotype, demonstrating that RsaM acts as a negative-feedback regulator that constrains QS activity after circuit activation. These findings redefine the role of RsaM in *P. fuscovaginae.* Rather than acting as a top-level repressor that prevents PfsI/R activation, RsaM serves as an intrinsic buffering mechanism that fine-tunes QS output and maintains signaling homeostasis once the system is activated. The mode of action of RsaM is currently unknown; however, it is likely that it takes place post-transcriptionally since the *rsaM* deletion in the mutant 0736IRIM did not result in a change in *pfsI* promoter activity. Interestingly, TofM in *B. glumae* BGR1 was recently reported to potentially bind the 5’UTR of the synthase *tofI* mRNA and block its translation. Unlike TofM, RsaM does not display significant homology to the RNA-binding protein RsmA (Goo & Hwang, 2023).

The regulatory role of RsaM has parallelism with RsaL, the built-in DNA-binding repressor of the LasI/R system in *P. aeruginosa* and of the PpuI/R system of *P. capeferrum* that counteracts LuxR-driven transcriptional activation of the synthase gene, thereby controlling the magnitude of the QS response (Rampioni et al., 2007). RsaL is found across *Pseudomonas* and *Burkholderia* species having LasI/R-like systems and is typically encoded by a gene embedded between the *lasI*-*lasR* gene pair. RsaL inactivation results in considerable overproduction of AHLs (Bertani & Venturi, 2004; Rampioni et al., 2007; Suárez-Moreno et al., 2008; Venturi et al., 2011).

QS can impose a substantial fitness burden on quorate populations, depending on the types and number of the target loci it regulates (Pai et al., 2012; Ruparell et al., 2016). The removal of repressive elements of the QS circuits often deregulates loci responsible for phenotypes that are timely and beneficial for bacterial populations in their ecological niches (Morici et al., 2007; Venturi et al., 2011; Gupta & Schuster, 2013). The high fitness cost of QS activation could have driven the evolution of a regulatory organization involving multi-layered repression and modulation. Given their role in virulence, it is probable that QS systems in *P. fuscovaginae* are triggered when the bacterial population encounters specific conditions in planta. The signals responsible for relieving this superimposed repression in *P. fuscovaginae*, and the regulatory pathways that convey these signals to the PfsI/R circuit, remain to be identified. We tested whether plant macerates could activate QS in *P. fuscovaginae* by monitoring AHL production and the activities of QS gene promoters. Under the conditions tested, plant macerates did not induce derepression of the QS circuitry (data not shown). Once this gatekeeping layer is removed, however, RsaM acts as an intrinsic buffering mechanism that opposes the self-amplifying PfsR positive-feedback loop. In doing so, RsaM likely maintains QS activity within an optimal range, preventing runaway activation and the potentially detrimental fitness consequences of uncontrolled signal production (Spacapan et al., 2023). A model of the regulatory organization of the PfsI/R circuit based on the results of this study is depicted in Figure 5.

**Figure 5.**
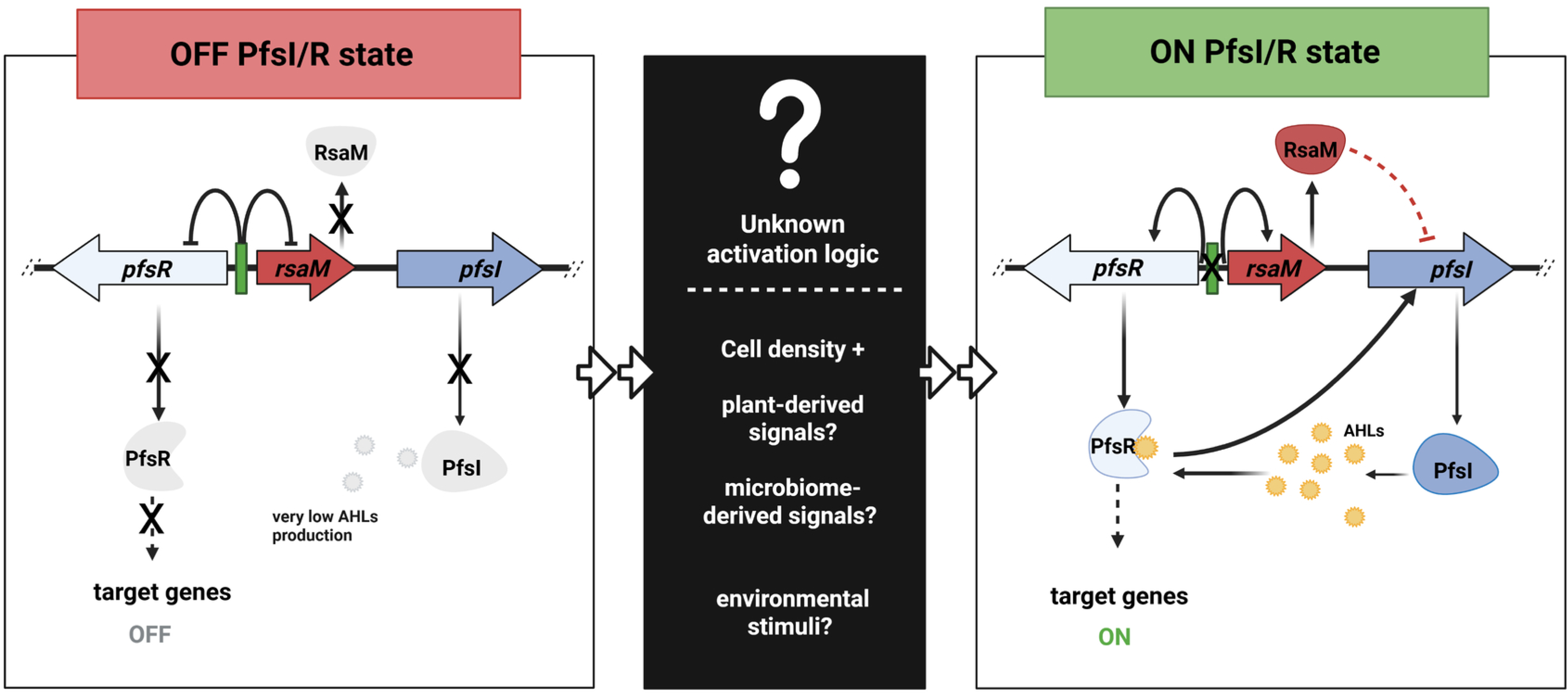
Current model of the regulatory architecture of the PfsI/R system.

## Acknowledgements

N.R. is a recipient of an ICGEB Arturo Falaschi Fellowship, and C.B. from the European Union’s Horizon Europe research and innovation programme under the Marie Skłodowska-Curie grant agreement No 101150379.

## Authors contributions

N.R. (Conceptualization, Data curation, Formal Analysis, Investigation, Methodology, Visualization, Writing – original draft, Writing – review & editing), I.B. (Project administration, Supervision), G.T. (Methodology, Validation), M.P.M. (Methodology, Data curation, Validation), C.B. (Conceptualization, Data curation, Investigation, Supervision, Validation, Visualization, Writing – original draft, Writing – review & editing), V.V. (Conceptualization, Funding acquisition, Investigation, Project administration, Resources, Supervision, Validation, Writing – original draft, Writing – review & editing).

